# SNP calling parameters have minimal impact on population structure and divergence time estimates for the rice blast fungus

**DOI:** 10.1101/2022.03.06.482794

**Authors:** Sergio M. Latorre, Thorsten Langner, Angus Malmgren, Joe Win, Sophien Kamoun, Hernán A. Burbano

**Affiliations:** Centre for Life’s Origins and Evolution, Department of Genetics, Evolution and Environment, University College London, London, United Kingdom; The Sainsbury Laboratory, University of East Anglia, Norwich, United Kingdom

**Keywords:** SNP calling, SNP filtering, fungi, plant pathogens, evolutionary rates, divergence times

## Abstract

**Objectives:** Accurate single-nucleotide polymorphisms (SNP) calls are crucial for robust evolutionary and population genetic inferences in genomic analyses. Such inferences can reveal the time-scales and processes associated with the emergence and spread of pandemic plant pathogens, such as the rice blast fungus *Magnaporthe oryzae* (Syn. *Pyricularia oryzae*). However, the specificity and sensitivity of SNP calls depend on the filtering parameters applied to the data. Here, we used a benchmarking approach to evaluate the impact of SNP calling on different population genetic analyses of the rice blast fungus, namely genetic clustering, topology of phylogenetic reconstructions and estimation of evolutionary rates.

**Results:** To benchmark SNP calling parameters, we generated a gold standard set of validated SNPs by sequencing nine *M. oryzae* genomes with both Illumina short-reads and Oxford Nanopore Technologies (ONT). We used the gold standard set of SNPs to identify the SNP calling parameter configuration that maximizes sensitivity and specificity. We found that the choice of parameter configurations can substantially change the number of ascertained SNPs, preferentially affecting SNPs segregating at low population frequency. However, SNP calling parameter configurations did not significantly affect the clustering of isolates in clonal lineages, the monophyly of each clonal lineage, and the estimation of evolutionary rates. We leverage the evolutionary rates obtained from each SNP calling parameter configuration to generate divergence time estimates that take into account the uncertainty associated with both the estimation of evolutionary rates and SNP calling. Our analysis indicates that *M. oryzae* clonal lineage expansions took place ~300 years ago.

## INTRODUCTION

Population genomics studies at the scale of hundreds of genomes are improving our understanding of the population structure, adaptive history and genome evolution of multiple plant pathogens [1]. This increasing availability of genome-wide genetic variation of fungal and other filamentous plant pathogens calls for a more rigorous and objective approach to the ascertainment of single nucleotide polymorphisms (SNPs) (henceforth SNP calling). Indeed, SNP calling parameters can substantially alter the conclusions reached by population genomic analyses. For instance, inferences that rely on low frequency SNPs, such as the estimation of evolutionary rates, are particularly prone to be affected by SNP calling strategies. Here, we use a benchmarking approach to investigate the impact of SNP calling parameters on population genomics analyses of the rice blast fungus *Magnaporthe oryzae*, arguably the most threatening fungal plant pathogen to world agriculture [2].

SNP calling is a multistep parameter-rich procedure that takes place prior to any evolutionary and phylogenetic inference. Therefore, a thorough exploration and careful selection of SNP calling parameters is crucial to obtain robust analytical results. A common practice to fine tune SNP calling parameters is to generate a set of validated - “gold standard” - SNPs. These SNPs are typically generated by independent sequencing platforms (e.g. [3]), which permits accounting for platform-specific error profiles and biases. The use of gold standard SNPs should then guide the choice of SNP calling parameters that minimize the error. However, even in the presence of a gold standard, establishing hard thresholds on SNP calling parameters is challenging, i.e. by looking at each parameter independently, a fraction of true SNPs will be removed, while a fraction of false SNPs will be kept. Alternatively, establishing dynamic thresholds is possible when high-quality SNPs are previously ascertained in thousands of individuals [4, 5]. With the exception of a few species such as model organisms and crops, the vast majority of species do not have genomic resources to enable dynamic thresholding. Thus, there is still a need for establishing optimal SNP calling approaches for organisms with relatively modest genomic resources, as with the majority of fungal plant pathogens.

This increasing availability of genome-wide genetic variation of fungal and other filamentous plant pathogens calls for a more rigorous and objective approach to SNP calling. This study was prompted by concerns about the specificity and sensitivity of SNP calling in genome analyses of the rice blast fungus *M. oryzae*. This fungus is widely distributed in rice growing regions throughout the world and the major biotic threat to rice production [2]. Recently, to document the global population history of *M. oryzae*, we analyzed a set of 131 publicly available genomes [6, 7]. This analysis revealed the presence of three globally distributed - pandemic - lineages of the rice blast fungus that arose in the last couple of hundred years [8].

To benchmark SNP calling in *M. oryzae*, we generated a gold standard variants dataset (GSVD) by sequencing a set of nine isolates with Illumina short-reads and Oxford Nanopore Technologies (ONT) [9]. We used the GSVD to benchmark different sets of filtering parameters by estimating their sensitivity and specificity, and their effect on the reconstruction of the population history of a group of 140 Illumina sequenced genomes [8, 9]. We found that among all the GATK summary statistics, Quality-by-Depth (QD) discriminates the best between true positives and true negatives. Consequently, we suggest that, even in the absence of a GSVD, SNPs can be accurately filtered based on the empirical distribution of QD values. Our analyses also revealed that the population structure of *M. oryzae* clonal lineages and the estimation of evolutionary rates are generally robust to different SNP filtering criteria. Moreover, by incorporating the uncertainty associated with both the calculation of evolutionary rates and SNP calling, we generated more realistic confidence intervals for dating the most recent common ancestor of the clonal lineages, which supports their expansion over the last ~300 years.

## MAIN TEXT

### Condensed Material and Methods

#### Datasets

To generate the gold standard variants dataset (GSVD) we used a set of nine rice-infecting *M. oryzae* isolates (henceforth benchmarking isolates) sequenced by using Illumina short reads (average depth: 23.5X; σ: 4.7) and assembled using sequencing reads generated with Oxford Nanopore Technologies (ONT) (Additional file 1: Table S1) [9]. We test different SNP calling parameters using a total of 140 rice-infecting *M. oryzae* genomes sequenced using Illumina short-reads. They include 131 previously published genomes [6, 7] that were jointly analyzed in Latorre *et al*., 2020 [8] and the above mentioned set of nine benchmarking isolates [9]. The general workflow of the analyses is presented in Additional file 1: Fig S1. A more detailed description of the methodology outlined below is presented in Additional file 1: Supplementary Methods.

#### Generation of the gold standard variants dataset (GSVD)

We ascertained SNPs in every benchmarking isolate independently for both the Illumina short reads and the ONT assemblies. The GSVD consisted of the union of sites that were concordant between the two technologies across all isolates. Similarly, we created a non-GSVD consisting in the union of discordant sites between the two technologies across all isolates (Additional file 1: Fig S2).

#### Selection of filtering parameters based on GATK summary statistics

To select filtering parameters based on GATK summary statistics, we first carried out a joint call of SNPs using the Illumina short reads from the nine benchmarking isolates. For the further analysis described in this section, we used the SNPs resulting from the joint call that were either present in the GSVD or in the non-GSVD. We then identified the summary statistics that significantly differ between the GSVD and non-GSVD. Since the distributions of GSVD and non-GSVD overlap for most summary statistics (Additional file 1: Fig S3), our main criterion to select filtering thresholds aimed at maximizing the fraction of GSVD and minimizing the fraction non-GSVD. We found that Quality-by-Depth (QD) was the summary statistic that better separates GSVD from non-GSVD (Additional file 1: Fig S4). Thus, we generated a dataset that optimizes this separation based on QD, henceforth “QD-based”. Additionally, we increase or decrease the number of SNPs by modifying the QD threshold creating a “Relaxed” and a “Stringent” dataset, respectively (Additional file 1: Supplementary Methods).

#### SNP filtering in a set of 140 Magnaporthe oryzae Illumina-sequenced genomes

To evaluate the effect of different filtering parameters we created a new joint call of SNPs that included a set of 131 *M. oryzae* isolates previously analyzed [8] and the nine benchmarking isolates that were used for the GSVD generation [9]. We filtered the SNPs resulting from the joint call of the 140 isolates using four different sets of parameters: Latorre *et al*., 2020 [8], Stringent, Relaxed and QD-based (Supplementary Table 2).

#### Detailed Material and Methods

See Additional file 1: Methods.

### Results and discussion

#### The Gold Standard Variant Dataset (GSVD) reveals the sensitivity and specificity of different SNP filtering criteria

To generate a benchmarking dataset, we used Illumina and ONT genome sequences of nine isolates of *M. oryzae* (Additional file 1: Table S1) [9]. To this end, we independently called SNPs from each of these genomes using a haploid model since field isolates of *M. oryzae* are haploid (Additional file 1: Supplementary Methods). Before identifying matching SNPs between the two sequencing platforms, we first compared the number of SNPs ascertained by each methodology. In the nine benchmarking isolates, we did not find significant differences in the number of SNPs called independently by Illumina and ONT (Wilcoxon signed rank test p > 0.05). On average, we found that ~85% of the positions covered by Illumina and ONT matched the reference genome allele, i.e. invariant sites. We used the union of concordant sites between technologies across all nine isolates to generate the GSVD, and the union of discordant sites between technologies across all nine isolates to generate the non-GSVD. The difference in the total number of SNPs between the GSVD (mean: 16,306) and non-GSVD (65,204) revealed that the non-GSVD is enriched with singletons that are likely sequencing errors, whereas GSVD likely harbors a high proportion of high confidence SNPs that were concordant between the two technologies (Additional file 1: Fig S2). Thus, we can use the properties of the SNPs present in the GSVD to evaluate different SNP filtering criteria.

To this purpose, we performed a joint call of SNPs of all nine benchmarking isolates using the Illumina data. In this SNP set, we then compared the distributions of six GATK summary statistics for SNPs overlapping the GSVD and non-GSVD (Additional file 1: Supplementary Methods). We found that five out of six GATK summary statistics distributions were different between GSVD and non-GSVD SNPs (Additional file 1: Fig S3). We used them to select cutoffs that optimized the separation between the GSVD and non-GSVD summary statistics distributions (Additional file 1: Supplementary Methods; Additional file 1: Fig S4).

Using the GSVD and non-GSVD we calculated the sensitivity (true positive rate) and specificity (true negative rate) of four different filtering criteria: Latorre *et al*., 2020 [8], Stringent, Relaxed and QD-based. Relative to Latorre *et al*., both the QD-based and the Relaxed filtering criteria increased the sensitivity in ~20% and the specificity in 1% and ~20%, respectively (Additional file 1: Fig S5). The Stringent filtering shows the highest specificity but the lowest sensitivity with less than 5% true positive rate. Our analysis showed that once a GSVD and non-GSVD are ascertained, QD is the summary statistic that discriminates the best between them, by maximizing GSVD and minimizing non-GSVD. Thus, in the absence of a validated set of variants, selecting SNPs around the median QD (~1 standard deviation unit in our case) will maximize GSVD and minimize non-GSVD (Additional file 1: Fig S4). While the rest of the analyzed GATK summary statistics did not discriminate GSVD from non-GSVD, their discriminatory power might be higher in cases where the overall sequencing quality of the datasets is lower, or when dealing with organisms with higher ploidy levels than *M. oryzae*, since the presence of heterozygous calls introduces further complexity to SNP callling.

#### SNP filtering criteria can substantially affect the number of SNP calls

The total number of SNPs varied substantially among different filtering criteria (Additional file 1: Fig S6A). Relatively to Latorre *et al.*, both QD-based and Relaxing filtering criteria increased the number of SNPs, while the Stringent filtering parameters reduced them. We found that both QD-based and Relaxed filtering criteria doubled the number of SNPs originally ascertained in Latorre *et al*. from about 48,000 to 100,000. About 70% of these SNPs were shared between the QD-based and Relaxed datasets, while almost all SNPs present in the Latorre *et al*. dataset were contained in the QD-based and Relaxed datasets (Additional file 1: Fig S6A). In contrast, the Stringent filtering parameters reduced the number of SNPs to ~50% of Latorre *et al*., which leads to a large number of uncalled SNPs. Since the effect of different SNP numbers on the population history of *M. oryzae* will depend on their distribution and frequencies among different genetic lineages. We sought to systematically quantify such effects in the next sections.

#### The population structure of *M. oryzae* clonal lineages is generally robust to different SNP filtering criteria

We evaluated the effect of different filtering criteria on the clustering of isolates in populations using both a Principal Component Analysis (PCA) based on pairwise Hamming distances (Fig 1) and a hierarchical clustering based on *f3*-outgroup statistics (Additional file 1: Fig S7). The PCA analysis of Latorre *et al*., QD-based and Relaxed filtering criteria showed comparable patterns, with similar percentages of the variance explained by the three first PCs in all three criteria (i.e., 71.51-73.11% for PC1). In contrast, in the Stringent filtering, PC1 alone explains 87.4% of the variance, which results in a lower percentage of the variance explained by other PCs and, subsequently, less structuring in the PCA. Such lack of structure was likely caused by the ascertainment used in the Stringent filtering and not by the number of SNPs, since the PCA structure was not affected when we randomly down-sampled Latorre *et al.*, Relaxed and QD-based SNPs to the size of Stringent (N = 7,702) (Additional file 1: Fig S8). The differences between the Stringent filtering and the three other criteria were also manifested when we analyzed the robustness of the clusters using Silhouette scores, a measure of how well samples cluster together, i.e. the higher the Silhouette score the better the clustering (Additional file 1: Fig S9). While Latorre *et al*., QD-based and Relaxed showed the highest average Silhouette scores when the number of clusters (*K*) was 4-5, for the stringent filtering the highest scores were found when *K*=2, showing again weak population structuring likely due to the low number of SNPs and the ascertainment bias of this filtering.

**Figure 1.**
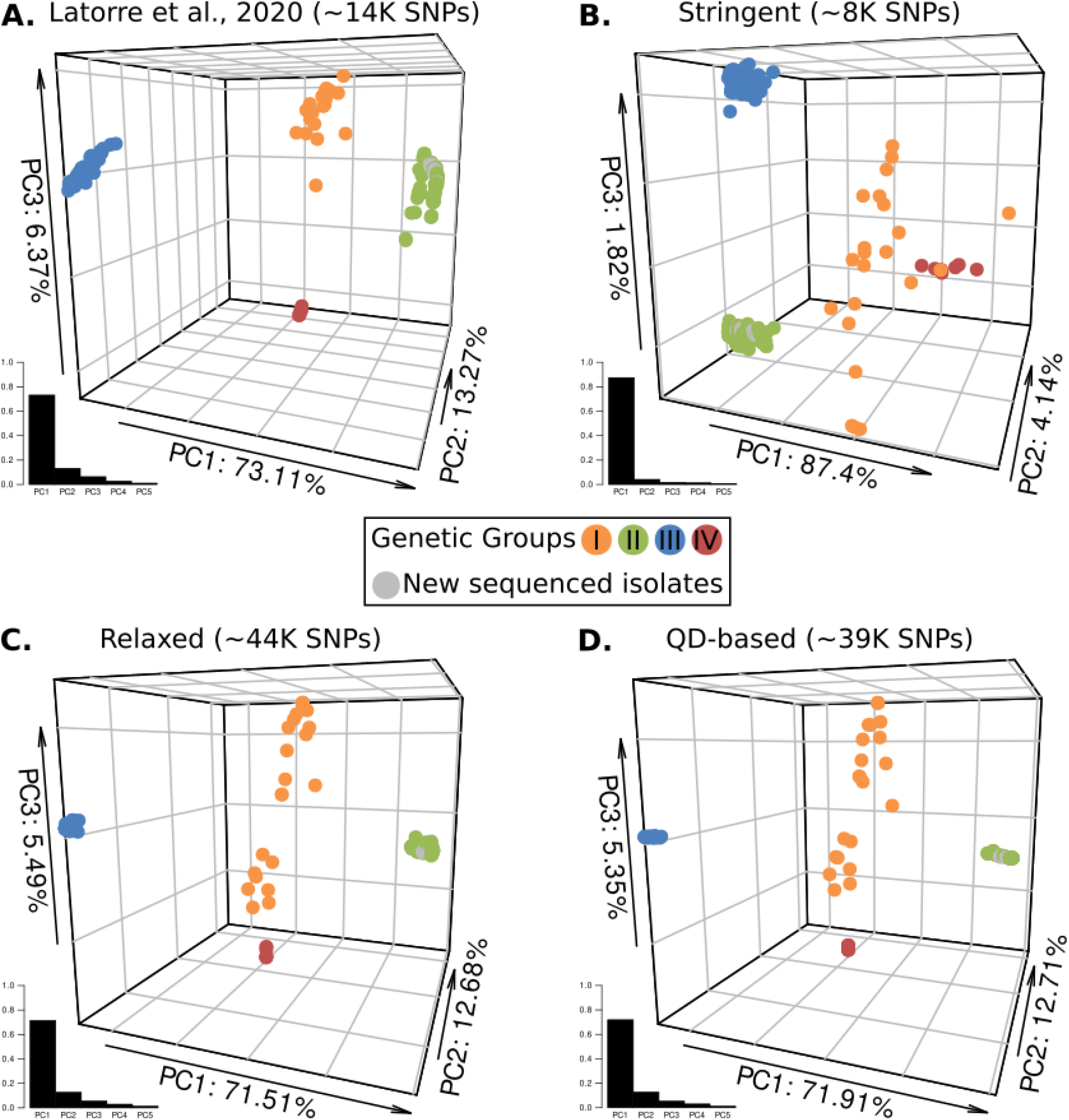
The population structure of clonal lineages II-IV is generally robust to different SNP filtering criteria. Each panel shows the separation of the fungal isolates in a 3-dimensional PCA space based on pairwise Hamming distances. The axes indicate the percentage of the total variance explained by each principal component (PC). We color-coded each isolate using the genetic classification established in Latorre *et al*, 2020 (Groups I-IV) [8]. The barplot adjacent to each 3D-PCA indicates the percentage of variance explained by the first ten PCs. The benchmarking isolates are depicted as gray points. In all panels the benchmarking isolates cluster tightly with isolates from group II (green points). The robustness of the clusters was assessed using Silhouette Scores (Additional file 1: Fig S9). **A.** Latorre et al. **B.** Stringent. **C.** Relaxed. **D.** QD-based. The number within parenthesis after each panel title refers to the total number of SNPs included in each dataset.

We have previously ascertained that the global population structure of the rice blast fungus consists of four groups (three clonal lineages and a genetically diverse group of isolates) [8]. Our per isolate analysis of Silhouette scores for the Latorre *et al*., QD-based and Relaxed criteria (all with 4 < K < 5), as well as the hierarchical clustering analysis (Additional file 1: Fig S7), showed that the three clonal lineages are consistently identified using these filtering criteria (Additional file 1: Fig S10). Moreover, incipient substructure within the diverse group of individuals could generate a possible fifth group (Fig 1C-D; Additional file 1: Fig S9-S10). We conclude that filtering criteria (i.e. QD-based, Relaxed) that double the number of SNPs originally ascertained in Latorre *et al*., provide more resolution to identify fine-grained population structure in the genetically diverse Group I. In summary, our analysis revealed the robustness of the three previously ascertained clonal lineages to SNP filtering criteria and the presence of fine-grained substructure within the genetically diverse Group I (orange).

#### SNP filtering parameters preferentially affects SNPs segregating at low frequency

Although the different filtering criteria resulted in the ascertainment of a wide range of total number of SNPs, the proportion of SNPs segregating in either the three clonal lineages (Group II-IV) or the genetically diverse Group I was maintained across all criteria (~25% and 75%, respectively) (Fig 2A). In a phylogeny, the maintenance of SNP proportions between genetic groups will probably result in an increase of branch lengths that remains proportional in all four genetic groups. The frequency of a given SNP in each clonal lineage corresponds to its placement in a phylogenetic context, i.e. singleton SNPs are placed on terminal branches, fixed SNPs are placed on the long internal branches that give rise to each clonal lineage, and SNPs that are neither singletons nor fixed are placed on internal branches within each clonal lineage (Figure 2C). Thus, to further explore the effect of the filtering criteria on the frequency of the ascertained SNP, we divided the SNPs of the four genetic groups into fixed, singletons or non-singletons (segregating at any frequency but not fixed). Whereas the three frequency-based categories of SNPs were present in the clonal lineages, fixed SNPs were almost absent in Group I, reflecting its high level of genetic diversity and heterogeneity (Fig 2B). The reduced occurrence of fixed SNPs in Group I, and the overall pattern of the frequency-based categories are not an artifact due to the uneven SNP number of each dataset, because the observed proportions remained similar when we downsampled each filtering criteria to the levels of the Stringent dataset (Additional file 1: Fig S11). However, we found that the two filtering criteria that increased the number of SNPs relative to Latorre *et al*., (QD-based and Relaxed), also increased the proportion of singletons in all genetic groups (Fig 2B). When placed in a phylogenetic context, singletons are located on terminal branches (tips of the trees) (Fig 2C), which may influence calculations that rely on SNPs segregating at low frequencies such as the estimation of evolutionary rates.

**Figure 2.**
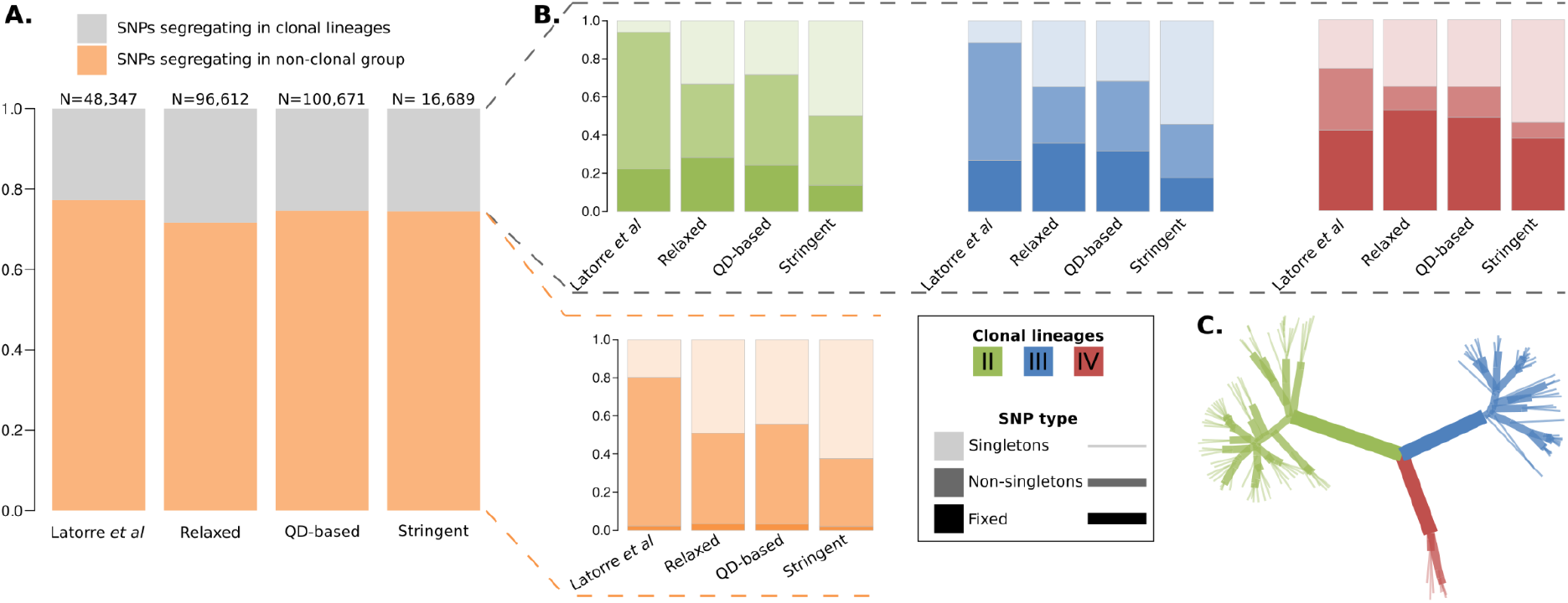
SNP filtering parameters affect preferentially SNPs segregating at low frequency. **A.** The stacked bars represent the fraction of SNPs segregating in all clonal lineages together and in the non-clonal diverse group for all SNP datasets. **B.** For each genetic group and for each SNP dataset the stacked bars display the fraction of SNPs that are singletons, fixed or at any intermediate frequency (non-singletons). The SNPs were classified in each of the frequency bins based on the numbers of individuals per genetic group carrying the non-reference allele. **C.** A schematic unrooted tree of clonal lineages II-IV indicating the distribution of every SNP type using the frequency bins described in **B.** Color hue and line thickness refer to different frequency-based SNP categories (inset).

#### SNP calling improves estimates of uncertainty in divergence time calculations

Given that different filtering parameters yield different numbers and qualities of SNP calls we reasoned that we could improve estimates of uncertainty in divergence time calculations. To investigate the effect of each filtering criteria, and their corresponding proportions of frequency-based categories on the estimation of evolutionary rates and the calculation of divergence times, we applied a Bayesian tip-dated phylogenetic analysis for each dataset. The number of new mutations in every isolate is expected to correlate with its sampling date, thus, such dates can be used to co-estimate the evolutionary rate with the time since the divergence of other internal nodes emerged [10]. Before performing the evolutionary rate analysis, it is crucial to test whether there is a phylogenetic temporal signal in the dataset (i.e. a positive correlation between sampling dates and genetic divergence). We found a phylogenetic temporal signal in all SNP datasets except for the Stringent filtering criteria (Additional file 1: Fig S12), which was the dataset with the lowest number of SNPs and lowest sensitivity (<5% true positive rate). Therefore, we discarded the Stringent dataset for the phylogenetic analysis. For the Bayesian phylogenetic analysis we used the sample collection dates for tip-calibration [8, 11] and restricted it to the three non-recombining clonal lineages. Due to its recombining nature, we excluded Group I, which was in fact the group which gained most SNPs when we applied the QD-based and relaxed filtering criteria. Thus, the vast majority of newly ascertained SNPs will not play any role in the estimation of evolutionary rates and calculation of divergence times.

We obtained a phylogenetic reconstruction that supports the monophyly of each clonal lineage with a posterior probability of one (Fig 3A). This topology supports previous analysis [8] and was also obtained for each filtering criteria independently (Additional file 1: Fig S13). In spite of the increased number of SNPs in the QD-based and Relaxed filtering criteria compared to Latorre *et al*., the three filtering criteria yielded evolutionary rates of the same order of magnitude that were not significantly different from each other (overlapping 95% Highest Posterior Density Intervals - HPD) (Fig 3B). The Bayesian phylogenetic analysis co-estimates *de novo* both the tree and the posterior distribution of the evolutionary rate and may have contributed to the robustness in the evolutionary rates estimations. The faster rates resulted in older divergence time estimations for the most recent common ancestor (TMRCA) for each clonal lineage. In contrast, the Relaxed filtering criteria generated both slower evolutionary rates in comparison to the rest (Fig 3B) as well as the widest confidence intervals for TMRCA of the different clonal lineages (Fig 3C). Moreover, the Effective Sample Size (ESS) values - a measure of the quality of the posterior distribution of a given parameter - reached with the Relaxed filtering criteria were consistently smaller for all the parameters (Table S4). Taken together, these results suggest a higher level of uncertainty on the estimations generated with the Relaxed filtering criteria in comparison to both Latorre *et al* and QD-based. The majority of differences between median estimates of TMRCA were smaller than 100 years, with only one median TMRCA of 157 years (clonal lineage IV) (Fig 3C; Additional file 1: Table S5). In addition to its oldest median divergence estimate, clonal lineage IV showed also the widest confidence interval and was the only clonal lineage divergence time that differs from our earlier estimates [8] (Additional file 1: Table S6), which could be due to its small sample size (N=7); the smaller the number of isolates/segregating SNPs, the lower the strength of the phylogenetic signal in this particular clonal lineage.

**Figure 3.**
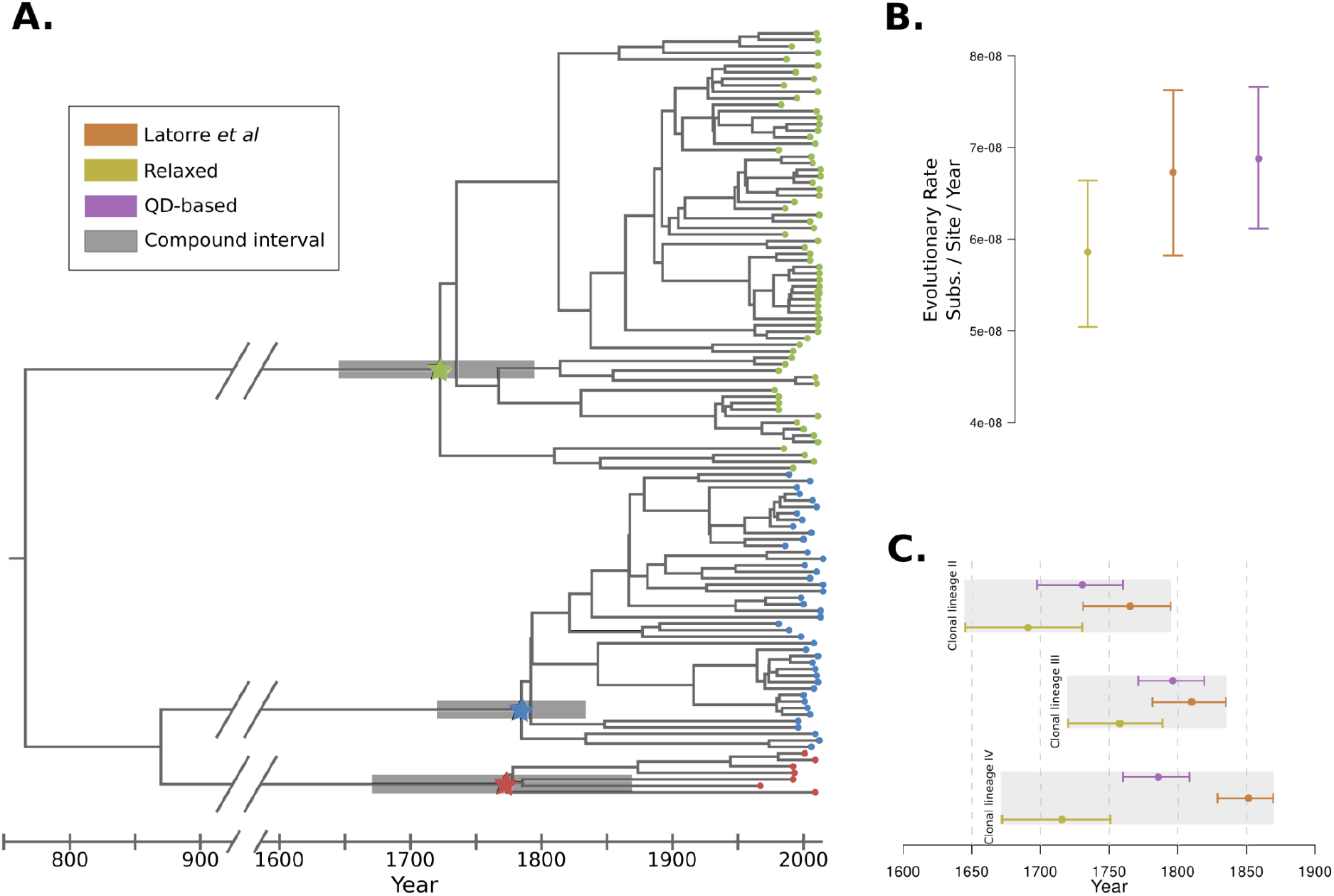
The use of different SNP filtering criteria improves the estimates of uncertainty of divergence time calculations. **A.** Bayesian tip-calibrated phylogenetic tree based on the combined posterior trees from the three SNP datasets. The stars indicate the most recent ancestor (MRCA) for clonal groups II (red), III (blue), and IV (red). The bars around the stars show the lowest and highest extremes from the 95% HPD for the divergence time estimate for the MRCA calculated in B. The inset shows the color code for each SNP dataset. **B.** Median and 95% Highest Posterior Density Intervals (HPD) intervals of the evolutionary rate for three SNP datasets. **C.** Divergence time estimations for the MRCA for each clonal lineage. The point indicates the median divergence time and the whiskers represent 95% HPD.

Our approach of varying SNP calling parameters enabled us to improve the estimates of uncertainty in the calculations of divergence times. To this end, we used the minimum and maximum upper and lower 95% HPD from the three filtering criteria to generate a compound TRMCA confidence interval (Fig 3A). In this way, in addition to the uncertainty on the TMRCA for each dataset derived from the posterior distribution of the evolutionary rate parameter, we incorporate the SNP uncertainty in the calculation of the TRMCAs. These calculations date the three clonal lineage expansions to ~300 years ago at most and are overall consistent with the previous calculations of Latorre et al [8].

## LIMITATIONS

We used a relatively small number of genomes sequenced by both Illumina and ONT to generate the Gold Standard Variants Dataset.

## List of abbreviations

*M. oryzae*: Magnaporthe oryzae
ONT: Oxford Nanopore Technologies
VQSR: Variant Quality Score Recalibration
GATK: Genome Analysis Toolkit
SNP: Single Nucleotide Polymorphism
GSVD: Gold Standard Variants Dataset
QD: Quality by Depth
REF: Reference
ALT: Alternative
ReadPosRankSum: Rank sum test for relative positioning REF versus ALT alleles within reads.
MQRankSum: Rank sum test for mapping qualities of REF versus ALT reads
BaseQRankSum: Rank sum test of REF versus ALT base quality scores
FS: Strand bias estimated using Fisher’s exact test
VCF: Variant Call Format
PCA: Principal Component Analysis
SFS: Site Frequency Spectrum
MCMC: Markov Chain Monte Carlo
MRCA: Most Recent Common Ancestor
TMRCA: Time to the Most Recent Common Ancestor
HPD: High Posterior Density
ESS: Estimated Sample Size
TMRCA: Time to the Most Recent Common Ancestor

## Declarations

### Ethics approval and consent to participate

Not applicable.

### Consent for publication

Not applicable.

### Availability of data and materials

The datasets and scripts generated during and/or analyzed during the current study are available in the Github repository, https://github.com/Burbano-Lab/rice-blast-variant-calling

### Authors’ contributions

SML, SK and HAB conceived the study. SML, AM, TL and JW designed and performed data analyses. SML, SK and HAB wrote the paper. All authors read and approved the final manuscript.

### Acknowledgements

We thank members of the Andrés, Burbano and Kamoun Laboratories and Sandra Álvarez-Carretero for useful discussions and input on data analysis; Aida Andrés and Michael Dannemann for comments on the manuscript; and Mark Farman for pointing out the stringency of the SNP calls performed in Latorre et al., 2020 [8].

## Additional file 1

### Supplementary Methods

#### Generation of the gold standard variants dataset (GSVD)

We ascertained SNPs in every sample independently for both the Illumina short reads and the ONT assemblies. We mapped the Illumina short reads to the GUY-11 reference genome (assembly number ASM236848v1; BioProject PRJNA354811) [12] using *bwa-mem* [13]. Subsequently, we identified the SNPs using the *GATK* haplotype caller [14]. For further analyses we did not use short insertions and deletions and only kept single nucleotide polymorphisms (SNPs). To ascertain the SNPs in the ONT dataset we used the genome assemblies reported by Win *et al*. [9] prior to the polishing step with Illumina short reads. In this way we can keep the independence of the two technologies in the ascertainment of variant sites. We mapped the unpolished ONT assemblies to the GUY-11 reference genome [12] using *minimap2* [15] and identified variant sites with *bcftools mpileup* [16]. We used mapping coverage as a proxy for mapping specificity and removed sites with a coverage depth greater than 1X. We only kept SNPs and filtered out insertions and deletions. For every isolate we identified concordant and discordant SNPs between the Illumina and the ONT datasets. The GSVD consisted of the union of concordant sites from all isolates. Similarly, we created a non-GSVD consisting in the union of discordant sites from all isolates (Additional file 1: Fig S2).

#### Selection of filtering parameters based on GATK summary statistics

To select filtering parameters based on GATK summary statistics, we first carried out a joint call of SNPs using the Illumina short reads from the benchmarking isolates. This step is necessary since the summary statistics of GATK based on per isolate callings are not directly translatable to the summary statistics resulting from jointly calling SNPs from a set of isolates. The joint call of SNPs was performed following GATK best practices [5] and the methodology described in Latorre *et al.,* 2020 [8]. For the further analysis described in this section we used only the SNPs resulting from the joint call that were either present in the GSVD or in the non-GSVD.

In order to identify the summary statistics that significantly differ between the GSVD and non-GSVD, we performed a Wilcoxon signed-rank test for every pair of summary statistics. Consequently, for further analysis we only used the following summary statistics: Quality by Depth (QD), rank sum test for relative positioning of reference (REF) versus alternative (ALT) alleles within reads (ReadPosRankSum), rank sum test for mapping qualities of REF versus ALT reads (MQRankSum), rank sum test of REF versus ALT base quality scores (BaseQRankSum), and strand bias estimated using Fisher’s exact test (FS). Since the distributions of GSVD and non-GSVD overlap for most summary statistics (Additional file 1: Fig S3), our main criterion to select filtering thresholds aimed at maximizing the fraction of GSVD and minimizing the fraction non-GSVD. We found that QD was the summary statistic that better separates GSVD from non-GSVD. Therefore, to set up upper and lower QD thresholds we identified the median of the QD GSVD distribution and calculated the magnitude of standard deviation units around the QD median that maximizes GSDV and minimizes non-GSDV. Henceforth, we refer to these SNPs as the QD-based set. Additionally, we used QD and the additional summary statistics to generate “Relaxed” and “Stringent” SNP datasets. The Relaxed set uses cutoffs that allow the inclusion of 90% of the GSVD without controlling for non-GSVD, whereas the Stringent set only allows the inclusion of 10% of non-GSVD without controlling for GSVD. Additionally, we used the filter set previously described in Latorre *et al.*, 2020 [8] with the aim to compare the effect of different filter sets on the robustness of previous inferences. A precise summary of the criteria used to generate each dataset is presented in (Additional file 1: Table S2). For each SNP dataset we defined true positives as the fraction of kept SNPs present in the GSVD and true negatives as the fraction of discarded SNPs present in the non-GSVD. Consequently, we defined false positives as the fraction of kept SNPs present in the non-GSVD and false negatives as the fraction of SNPs present in the GSVD and discarded in the filtered set (Additional file 1: Fig S4).

#### SNP filtering in a set of 140 *M. oryzae* Illumina-sequenced genomes

Initially, we performed the joint call of SNPs for 140 *M.oryzae* isolates following *GATK* best practices [5] as described in Latorre *et al.,* 2020 [8]. The joint call of SNPs included a set of 131 *M. oryzae* isolates previously analyzed [8] and the benchmarking isolates that were used for the GSVD generation [9]. We filtered the *vcf* file resulting from the joint call of the 140 isolates using four different sets of parameters: Latorre *et al*., 2020, stringent, relaxed and QD-based (Additional file 1: Table S2). Additionally, for analyses in which an outgroup was required, we included the wheat-infecting blast fungus isolate BTJP-4(12) [17], and filtered the SNP using the four previously described criteria. The type of dataset used on each analysis and its corresponding number of SNPs is shown in Additional file 1: Table S3. To detect SNPs intersecting among the filtered datasets we used the *R* package *UpSetR* [18] (Additional file 1: Fig S6).

#### Clustering of fungal isolates in populations

To evaluate the effect of the filters on the number of inferred population clusters, we used two different approaches. In both cases, we did not allow for missing data and used SNPs with full-information in all isolates. First, we performed a PCA based on pairwise Hamming distances (Fig 2) and retained the information from the first three components as they explained at least 90% of the total variance. To identify population clusters we computed Silhouette Scores [19, 20] with a number of clusters (*K*-values) ranging between 2 and 7. We selected the optimal *K*-value based on the highest average of Silhouette Scores. To evaluate the robustness of the population clusters we re-sampled isolates with replacement 1000 times, recalculated Silhouette Scores and generated bootstrap confidence intervals (Additional file 1: Fig S9).

We also used *f3*-outgroup statistics [21] to calculate distances for the configuration *f3*(A, B; outgroup) where A and B are all possible 9,730 pairwise comparisons and outgroup correspond to the wheat-infecting isolate. The computations were performed using *qp3Pop* from *AdmixTools* [22]. The distances were clustered with a Hierarchical Clustering algorithm using the *hclust* function in *R* (Additional file 1: Fig S7).

#### Effect of filtering criteria on the proportion of SNPs segregating at different frequency categories

To analyze the impact of the different filtering parameters on the frequency of the ascertained SNPs, we divided the SNPs segregating in each genetic group based on their within-genetic-group frequency, namely fixed, singleton or non-singleton (non-fixed SNPs present in more than one isolate). The inventory and filtering of SNPs based on their frequency was carried out using *bcftools* [16].

#### Phylogenetic analysis and calculation of evolutionary rates

We sought to analyze the impact of the different filtering parameters on the robustness of the phylogenetic reconstructions and the estimation of evolutionary rates of clonal lineages of the rice blast fungus (genetic groups II, III and IV). We first tested for the presence of a temporal signal on each dataset by correlating phylogenetic distances with collection dates. We used *RAxML-ng* [23] to generate Maximum Likelihood phylogenies from every filtering criteria dataset with 1,000 bootstrap replicates. For each tree, we then estimated the midpoint root and all root-to-tip distances using the *Biopython* library [24]. We calculated *Pearson’s r* correlation coefficients between the root-to-tip distances and the collection dates. Moreover, we measured the robustness of the correlations by sampling with replacement (1,000 iterations). To test whether the obtained correlation was greater than expected by chance, we randomly permuted the distances and the collection dates (1,000 iterations) (Additional file 1: Fig S12). Due to the absence of a phylogenetic signal on the Stringent filtering criteria, we excluded this SNP dataset for further analyses.

Using concatenated genome-wide SNPs, we performed a Bayesian phylogenetic reconstruction with BEAST2 (v.2.6.3) [11, 25] through the CIPRES gateway [26]. As described in Latorre *et al.*, 2020 [8], we carried out the analysis using the collection dates as prior information to build a tip-dated phylogeny. We chose a Coalescent Extended Bayesian Skyline [27] approach to reduce the effect of demographic history assumptions.

To determine the number of invariant sites to be included in the analysis of each SNP dataset, we first estimated the total number of base pairs of the *M. oryzae* reference genome covered by Illumina reads in all isolates, i.e. the intersection of covered positions in all isolates (Total Covered Genome). Afterwards, we ascertained the number of invariant sites by subtracting from the Total Covered Genome the total number of SNPs that passed the filtering criteria for each dataset (Additional file 1: Table S4). We assumed that the SNPs that did not pass the filtering were invariant sites, thus they were not included in the subtraction (Table S3). We included the total number of invariant sites as a “constantSiteWeights” tag in the *BEAST2-XML* input file. We reported results based on the merged posterior probability values of four independent MCMC chains per SNP dataset (Additional file 1: Table S5). The length of each chain was 10 million iterations.

### Data availability

All custom scripts used in this work as well as the pipeline details can be found at: https://github.com/Burbano-Lab/rice-blast-variant-calling

---

**Figure S1.**
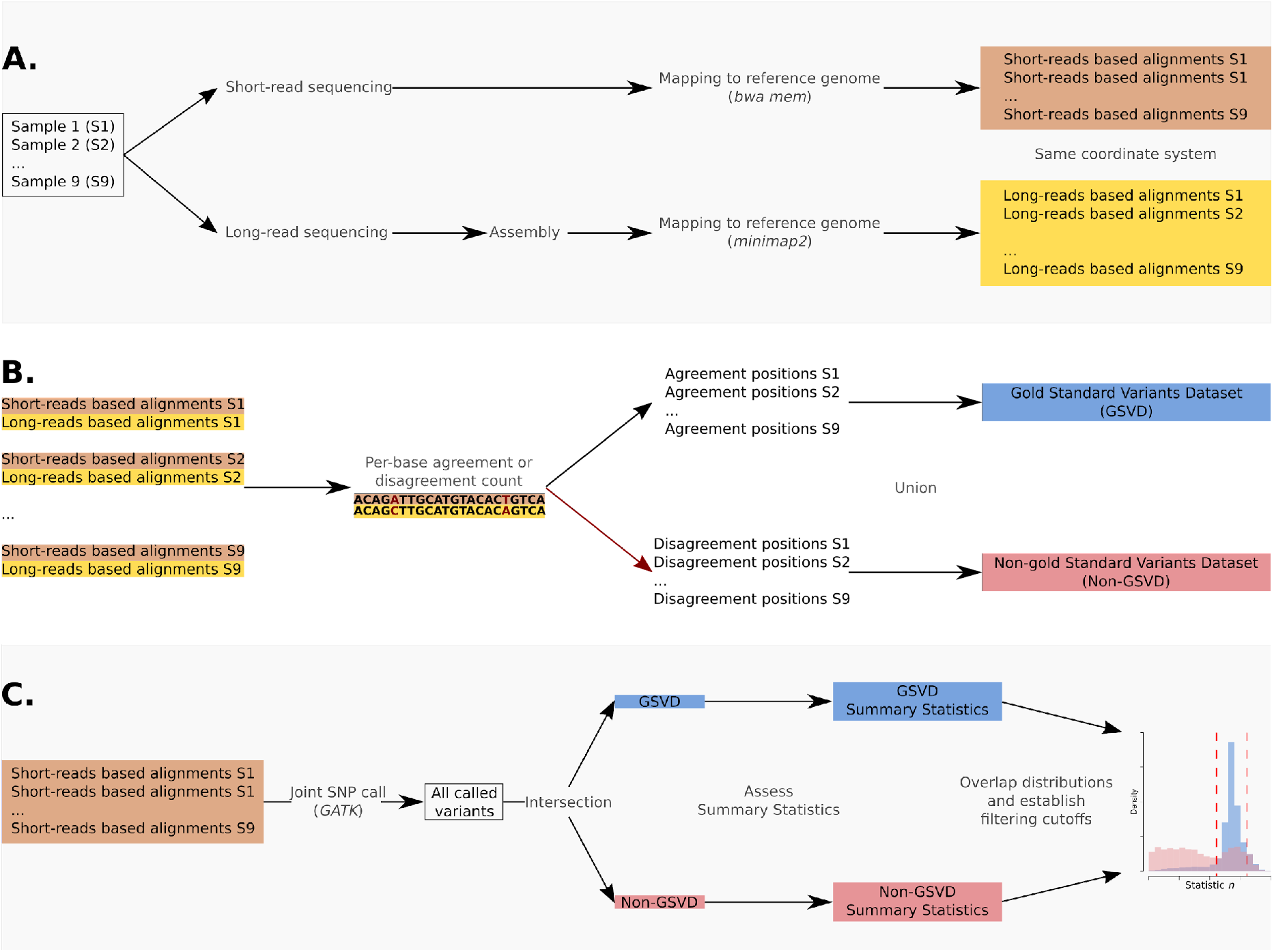
General workflow. **A.** Each sample is sequenced with both Illumina and and Oxford Nanopore Technologies (ONT) and mapped to the same reference genome. **B.** A Gold Standard Variant Dataset (GSVD) and a non-GSVD are the result of the union of the agreement and disagreement of the SNP calls of both technologies, respectively. **C.** Various distributions of summary statistics are extracted from a joint SNP call using Illumina short-reads. The intersection of these joint SNP calls and both GSVD and non-GSVD is used to calculate hard filtering cutoffs.

**Figure S2.**
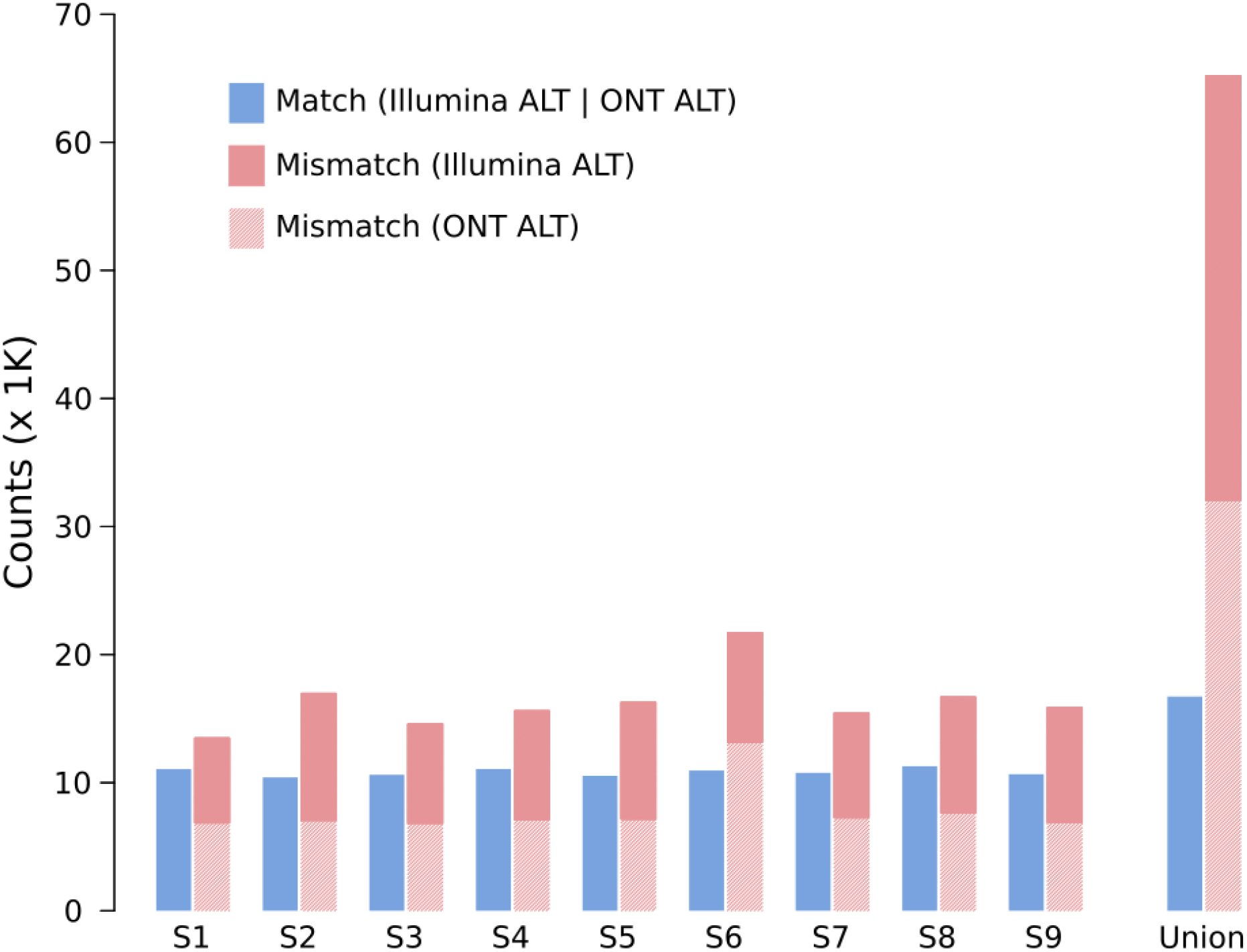
Concordance and discordance of SNPs identified using Illumina short reads and Oxford Nanopore Technology (ONT) long reads. Each pair of bars represents the proportion of non-reference alleles called when both methods agree (blue) or disagree (red) for each of the benchmarking isolates (S1 to S9). The last pair of bars represents the combined counts for all samples.

**Figure S3.**
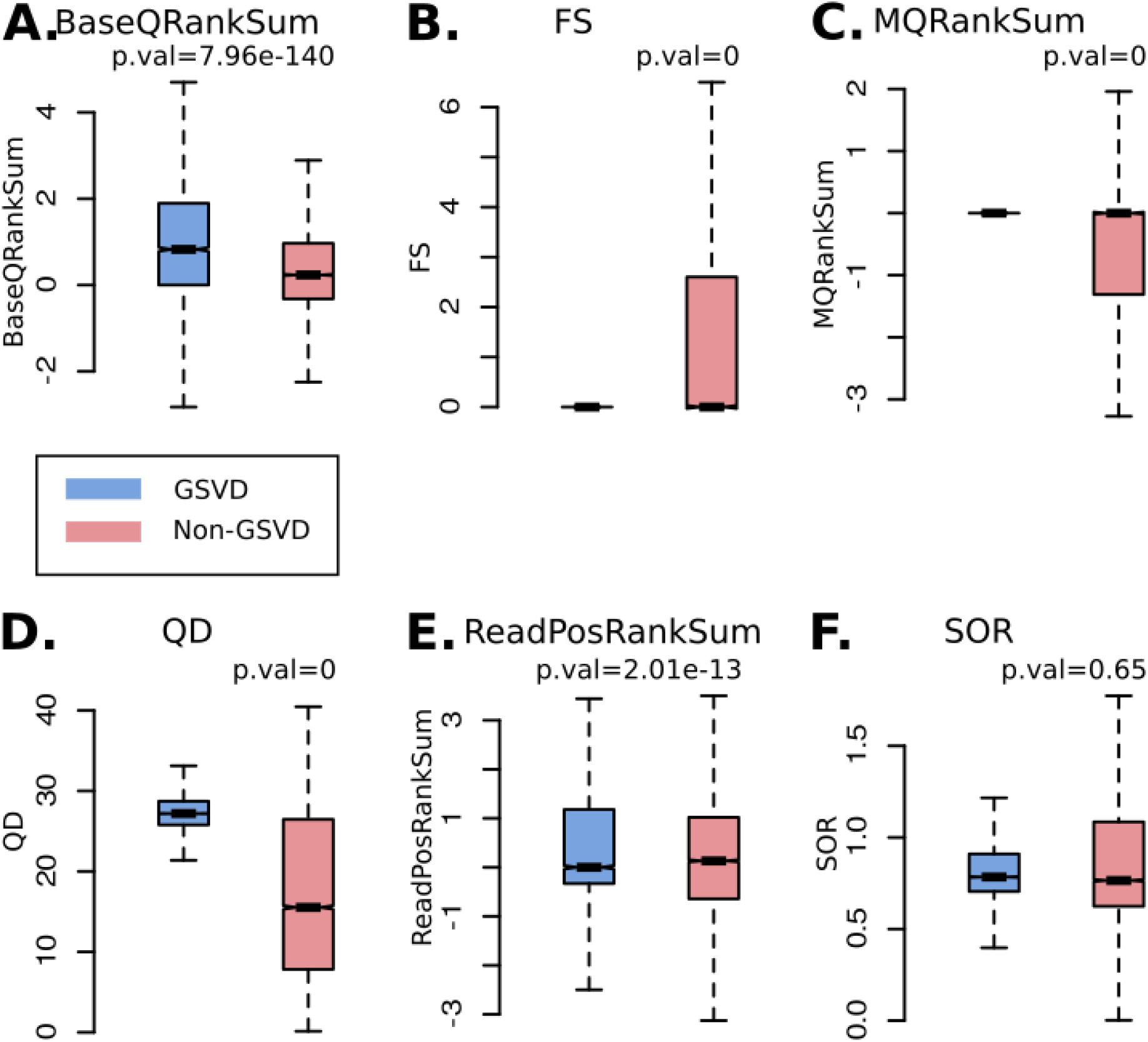
Comparison between the Gold Standard Variants Dataset (GSVD) and non-GSVD for different summary statistics calculated using the Genome Analysis Toolkit (GATK). Each boxplot shows the distribution of different summary statistics for GSVD (blue) and non-GSVD (red) as notched boxplots: **A.** Rank sum test of REF versus ALT base quality scores (*BaseQRankSum*), **B.** strand bias estimated using Fisher’s exact test (*FS*), **C.** rank sum test for mapping qualities of REF versus ALT reads (*MQRankSum*), *D.* quality by depth (*QD*), *E.* rank sum test for relative positioning of reference (REF) versus alternative (ALT) alleles within reads (*ReadPosRankSum*), **F.** strand odds ratio (*SOR*). Wilcoxon signed-rank test *p values* are shown for each paired comparison.

**Figure S4.**
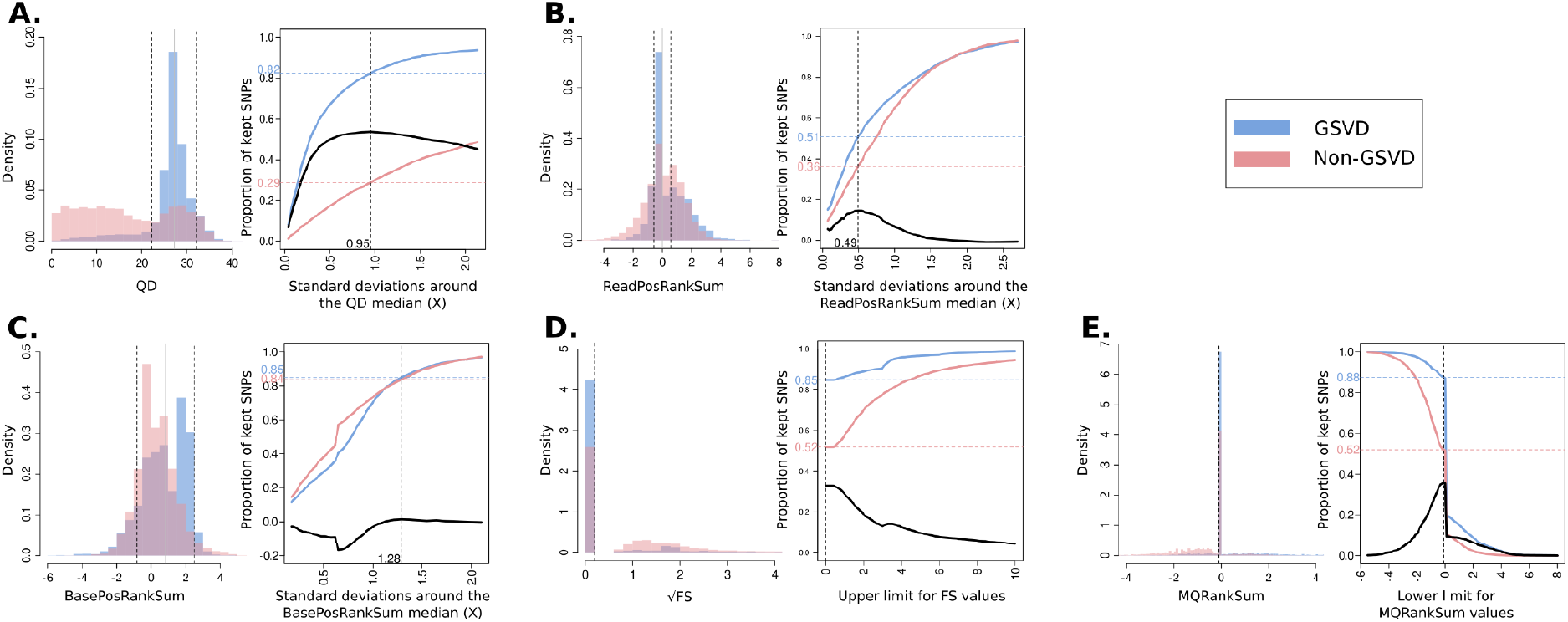
Selection of filtering cutoffs based on summary statistics from the Genome Analysis Toolkit (GATK). The distributions for each summary statistic are shown as histograms and the proportions of SNPs kept given a particular cutoff are shown in the adjacent plots. **A.** Quality by depth (*QD*), **B.** rank sum test for relative positioning of reference (REF) versus alternative (ALT) alleles within reads (*ReadPosRankSum*), **C.** rank sum test of REF versus ALT base quality scores (*BaseQRankSum*), **D.** strand bias estimated using Fisher’s exact test (*FS*), **E.** rank sum test for mapping qualities of REF versus ALT reads (*MQRankSum*). In all instances, blue colors represent GSVD and red colors non-GSVD. Vertical dotted black lines delimit the cutoffs for which the proportion of GSVD is maximized while the proportion of non-GSVD is minimized. **A.**, **B.**, and **C.**, are bounded with lower and upper limits while **D.** and **E.** are bounded only by the upper limit. In all cases, black solid lines represent the difference between GSVD and non-GSVD proportions.

**Figure S5.**
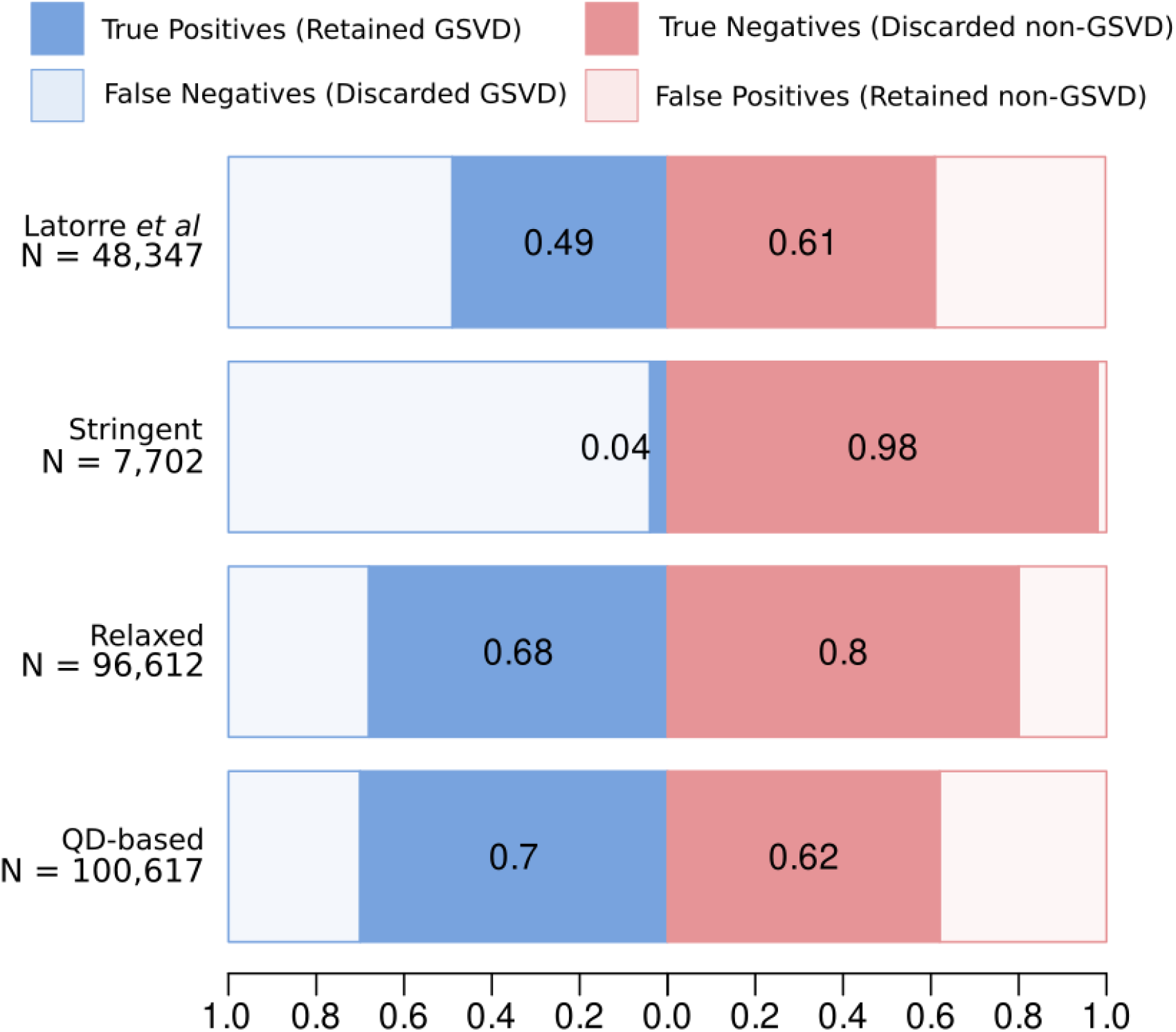
Sensitivity and specificity of different SNP filtering criteria. Each pair of horizontal stacked barplots shows the sensitivity (dark blue) and specificity (dark red) of each SNP datasets as proportions. Total SNP numbers are shown.

**Figure S6.**
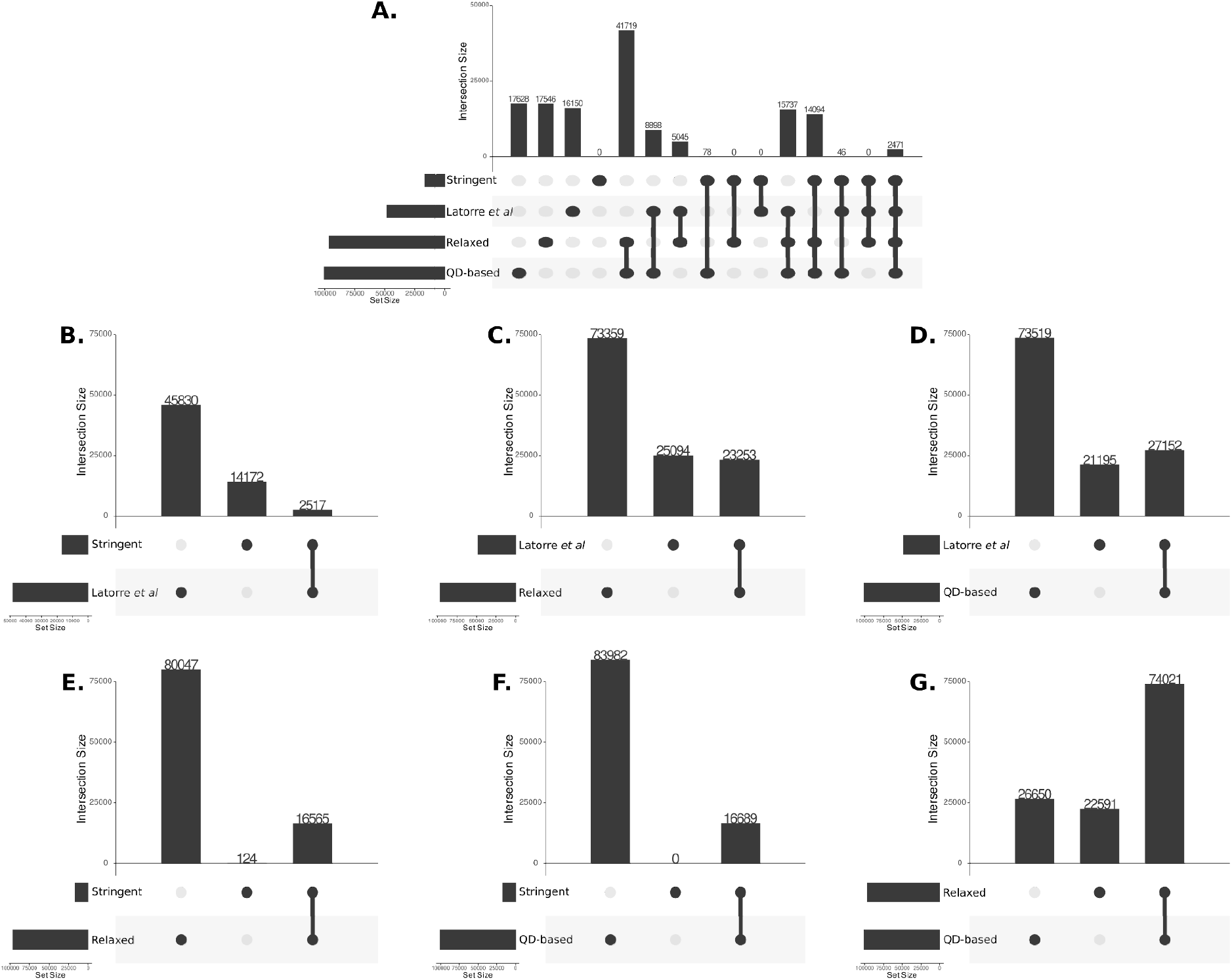
Intersection of SNPs among filtered datasets. The panels below each set of bars represent different intersection configurations. Multiple dot arrays (dots linked by lines) represent intersections, while single dots represent the private SNPs for the correspondent SNP datasets. The height of the vertical bars corresponds to the number of SNP for all possible intersections (**A**) and all pairwise intersections (**B-G**).

**Figure S7.**
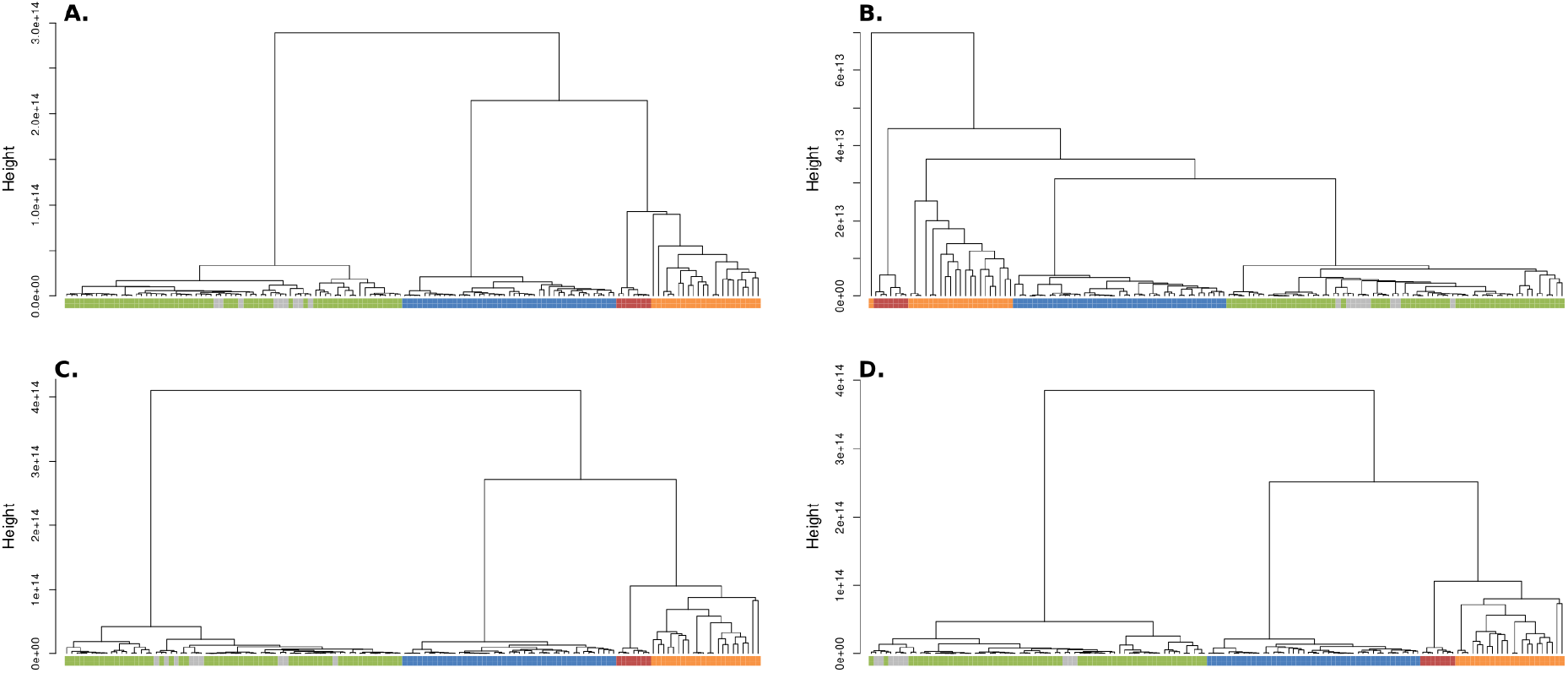
*f3*-outgroup based hierarchical clusters per filtered SNP dataset. The panels show the different hierarchical cluster based dendrograms calculated from pairwise *f3*-outgroup distances: **A.** Latorre *et al*, **B.** Stringent, **C.** Relaxed, **D.** QD-based. The colors correspond to those in Latorre *et al*, 2020 [8].

**Figure S8.**
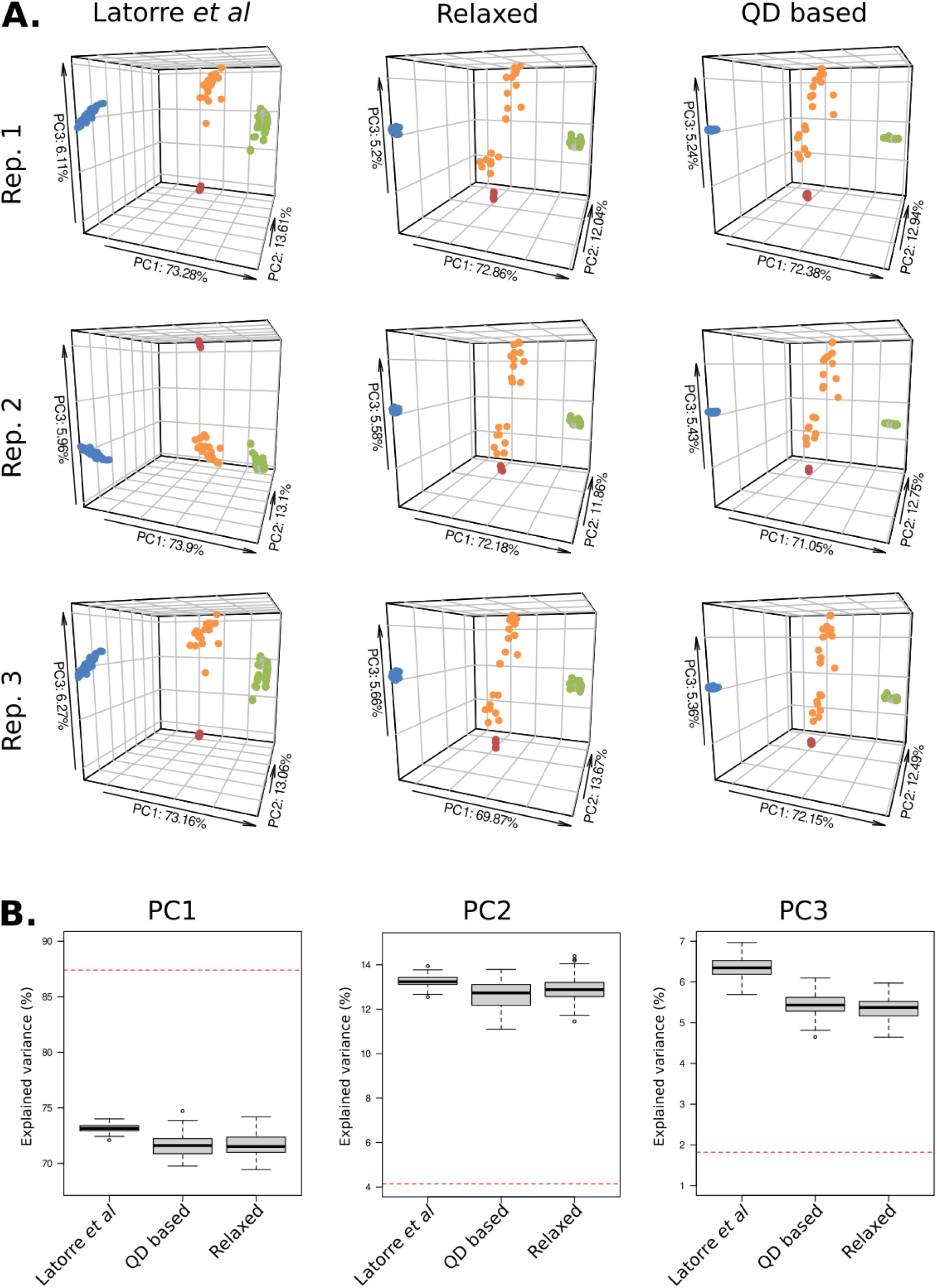
Robustness of Principal Component Analysis. **A.** PCA analyses of Latorre *et al*, Relaxed and QD-based SNP datasets after each set was randomly down-sampled to the size of the Stringent filtering set (N = 7,702) (three replicates shown). **B.** Boxplots represent the distributions of the percentage of explained variance for the three first principal components after 100 iterations of random downsampling. The horizontal red dotted line corresponds to the percentage of explained variance for the Stringent SNP set.

**Figure S9.**
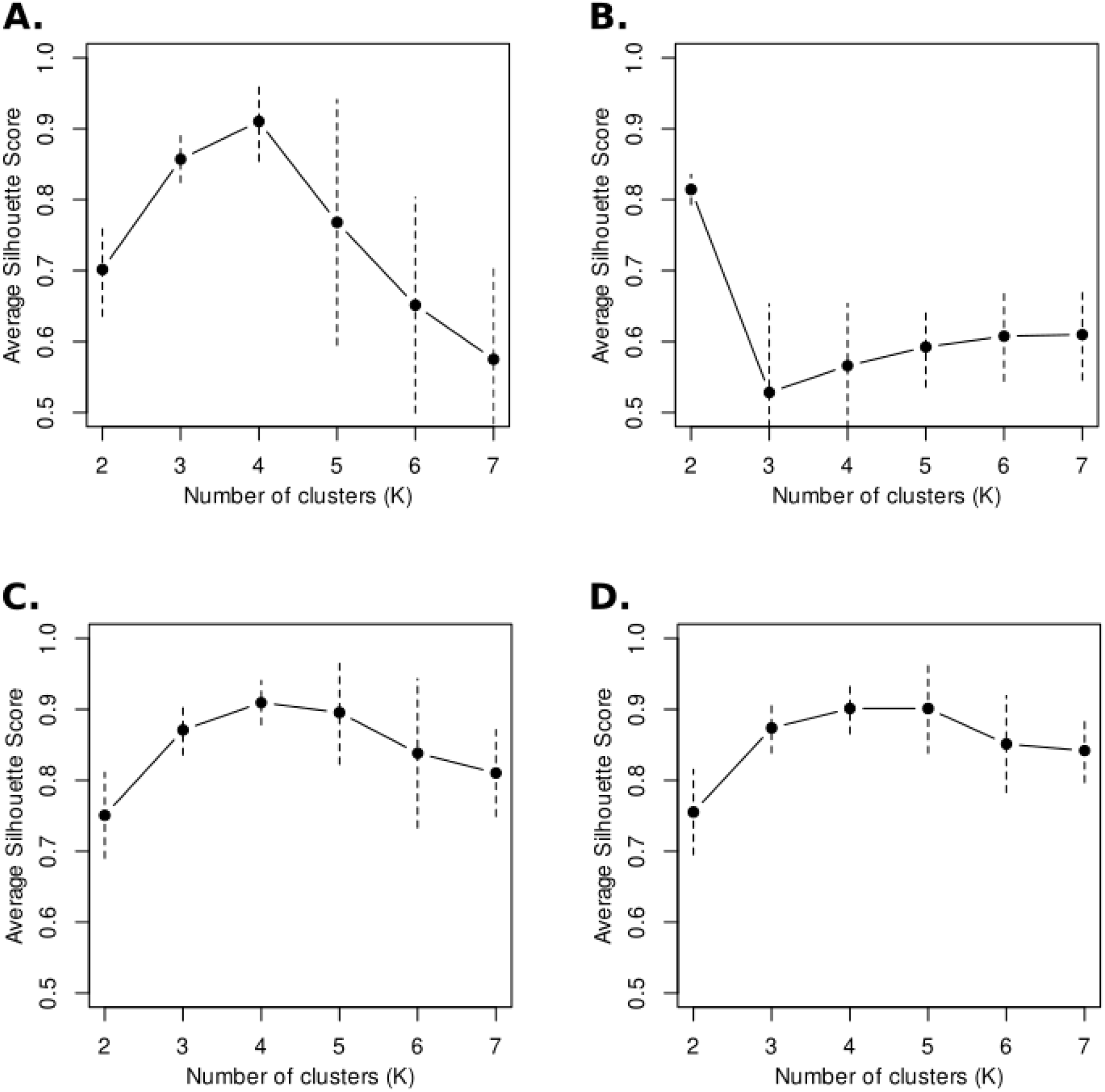
Silhouette Scores with bootstrap. Each panel represents the average Silhouette Scores based on the coordinates of the 3 firsts PC’s of each filtered SNP dataset (Fig 1) (**A.** Latorre *et al.*, **B.** Stringent, **C.** Relaxed, **D.** QD-based) when the number of clusters (*K*) range from 2 to 7. The dots correspond to the median value of 1000 silhouette scores computed by re-sampling isolates with replacement. The vertical dotted lines represent 95% confidence intervals.

**Figure S10.**
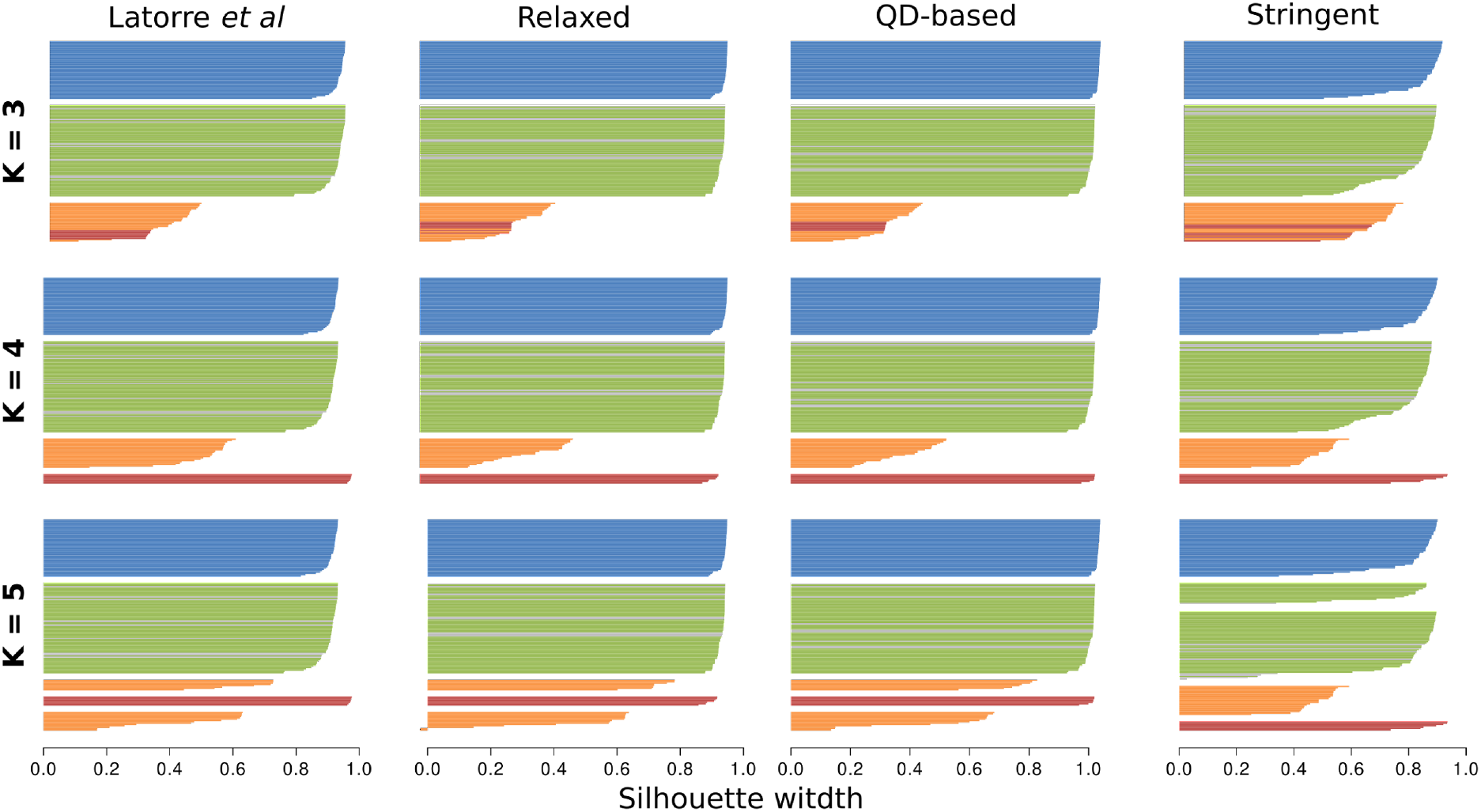
Per-isolate Silhouette Scores. Silhouette scores were computed based on the coordinates of the 3 firsts PC’s of each filtered SNP dataset (columns) when the number of clusters (*K*) range between 3 and 5 (rows). Each bar represents the silhouette score per isolate and each cluster of bars correspond to the identity cluster. Individuals were colored according to Latorre *et al,* 2020 [8], and the benchmarking isolates are colored in gray.

**Figure S11.**
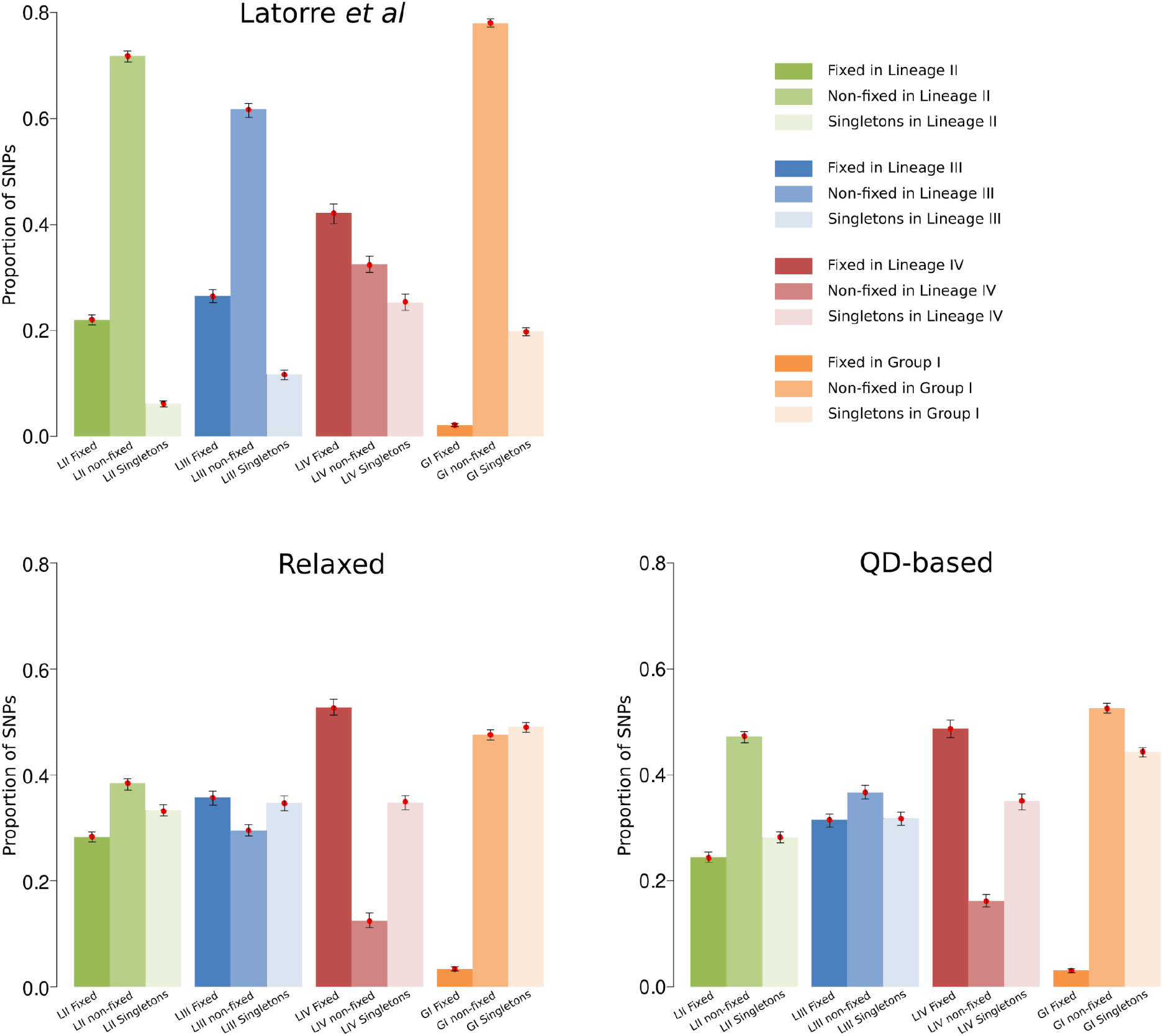
Effect of filtering criteria on the proportion of SNPs segregating at different frequency categories. SNP datasets from Latorre *et al*., Relaxed and QD-based were randomly downsampled to the size of the Stringent SNP set (N = 16,689). The height of each bar corresponds to the mean value and the whiskers represent the 2.5% and 97.5% quantiles of the distribution. Different color hues depict the three frequency categories (fixed, non-fixed and singletons) for each genetic group. The red dots represent the calculated values with the complete SNP datasets per filtering criteria as in Fig 2B.

**Figure S12.**
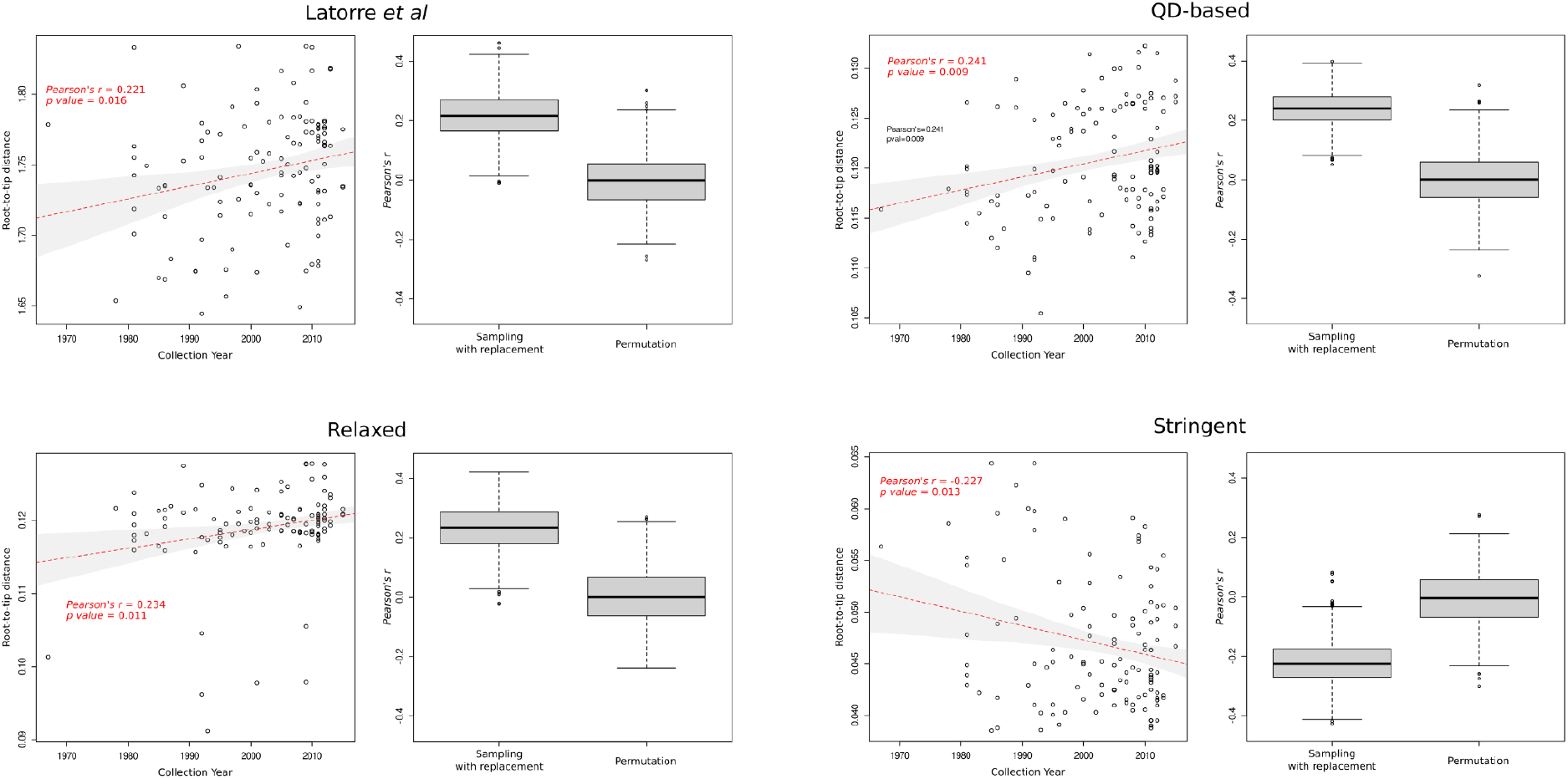
Effect of SNP filtering parameters on temporal signal of tip-calibrated phylogenies. The scatterplots show the correlation between root-to-tip distances from Maximum-Likelihood phylogenies calculated under the different filtering criteria against the collection years of the isolates. The red dotted lines depict the associated linear regressions together with their respective significance values in red colors. The boxplots show the distributions of the *Pearson’s r* correlation coefficient of 1,000 replicates of sampling with replacement and permutations of the root-to-tip distances and the collection years, respectively.

**Figure S13.**
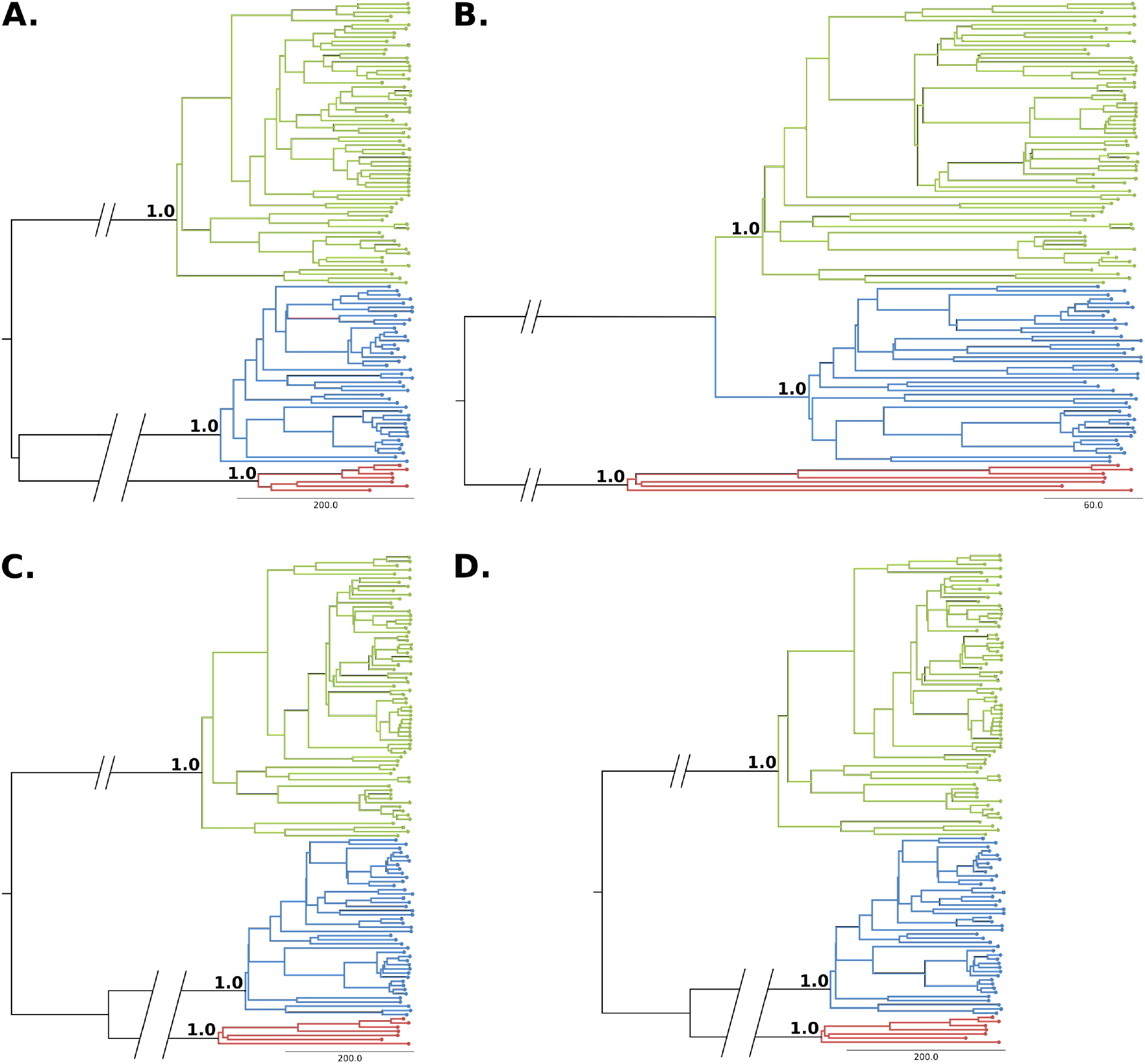
Bayesian phylogenies. Maximum credibility midpoint rooted trees calculated for each filtered SNP dataset. **A.** Latorre *et al*. **B.** Stringent. **C.** Relaxed. **D.** QD-based. The tree branches were colored according to Latorre et al, 2020 [8]. The values on the MRCA nodes for each clonal lineage indicate the posterior probability support.

**Table S1.** European Nucleotide Archive (ENA) accession numbers of the newly sequenced isolates [9].

| **Isolate** | **Illumina Short Reads** | **Oxford Nanopore Long Read Assemblies** |
| --- | --- | --- |
| **AG002** | ERR4757123 | GCA_905067045.1 |
| **AG038** | ERR4757126 | GCA_905067005.1 |
| **AG059** | ERR4757128 | GCA_905066965.1 |
| **AG098** | ERR4757129 | GCA_905067015.1 |
| **San_Andrea** | ERR4757131 | GCA_905067085.1 |
| **AG006** | ERR4757124 | GCA_905067025.1 |
| **AG032** | ERR4757125 | GCA_905067055.1 |
| **AG039** | ERR4757127 | GCA_905067035.1 |
| **PR003** | ERR4757130 | GCA_905067075.1 |

**Table S1.** Description of filters.

| Statistic \ Filter Set | Latorre *et al* | Relaxed | Stringent | QD-based |
| --- | --- | --- | --- | --- |
| **QD** | $X < 5.0$ | $X < 19.72884$<br>$X > 34.65116$ | $X < 25.71431$<br>$X > 28.66569$ | $X < 22.21431$<br>$X > 32.16569$ |
| **ReadPosRankSum** | $X < -2.0$<br>$X > 2.0$ | $X < -2.234249$<br>$X > 2.234249$ | $X < -0.0971292$<br>$X > 0.0971292$ | NA |
| **FS** | NA | $X > 2.6$ | $X > 0.1$ | NA |
| **MQRankSum** | $X < -2.0$<br>$X > 2.0$ | $X < -0.5$<br>$X > 4.0$ | NA | NA |
| **QUAL** | $X < 5,000$ | NA | NA | NA |
| **BaseQRankSum** | $X < -2.0$<br>$X > 2.0$ | NA | NA | NA |
| **MQ** | $X < 20.0$ | NA | NA | NA |

**Table S2.** Type of analyses and number of SNP per dataset.

| SNP dataset | Analyses | Filter |  |  |  |
| --- | --- | --- | --- | --- | --- |
|  |  | Latorre *et al* | Stringent | Relaxed | QD-based |
| Rice-infecting isolates. All SNPs | QC | 48,347 | 16,689 | 96,612 | 100,671 |
| Rice-infecting isolates. Full information | PCA-based population structure analyses | 13,637 | 7,702 | 43,688 | 39,139 |
| Rice-infecting isolates with outgroup. All SNPs |  | 293,259 | 83,376 | 357,548 | 323,191 |
| Rice-infecting isolates with outgroup. Full information | *f*3-outgroup-based population structure analyses | 194,463 | 51,986 | 216,204 | 182,661 |
| Clonal rice-infecting isolates. Full information | SVD-quartet based phylogeny. Tip calibrated bayesian phylogeny | 11,013 | 4,259 | 27,318 | 25,546 |

**Table S3.** Invariant sites calculation summary

| **Filter** | **Total**<br>**Covered**<br>**Genome** | **Invariant sites** | **A** | **C** | **G** | **T** |
| --- | --- | --- | --- | --- | --- | --- |
| Relaxed | 32'770.348 | 32'743.087 | 7'867.377 | 8'502.216 | 8'515.627 | 7'857.867 |
| Latorre |  | 32'759.382 | 7'869.949 | 8'507.698 | 8'521.249 | 7'860.486 |
| QD |  | 32'744.921 | 7'867.626 | 8'502.807 | 8'516.305 | 7'858.183 |

**Table S4.** Posterior summary statistics for Bayesian tip-calibrated phylogenies

| #Evolutionary_clock_rate |  |  |  |  |  |  |  |  |  |
| --- | --- | --- | --- | --- | --- | --- | --- | --- | --- |
|  | Mean | Stderr of mean | Stdev | Variance | Median | Geom. mean | Lower 95HPD | Upper 95HPD | ESS |
| Relaxed | 5.85E-08 | 3.29E-10 | 4.07E-09 | 1.66E-17 | 5.86E-08 | 5.84E-08 | 5.04E-08 | 6.62E-08 | 153.1275 |
| Latorre | 6.74E-08 | 3.45E-10 | 4.65E-09 | 2.16E-17 | 6.74E-08 | 6.72E-08 | 5.85E-08 | 7.66E-08 | 181.3641 |
| QD | 6.88E-08 | 2.10E-10 | 3.96E-09 | 1.57E-17 | 6.88E-08 | 6.87E-08 | 6.11E-08 | 7.66E-08 | 356.5619 |
| #Divergence_estimation_Clonal_Lineage_II |  |  |  |  |  |  |  |  |  |
|  | Mean | Stderr of mean | Stdev | Variance | Median | Geom. mean | Lower 95HPD | Upper 95HPD | ESS |
| Relaxed | 1688.8472 | 1.7913 | 22.112 | 488.942 | 1690.6871 | 1688.7018 | 1645.1194 | 1730.0967 | 152.3773 |
| Latorre | 1763.8986 | 1.3422 | 16.8131 | 282.6797 | 1765.1675 | 1763.8183 | 1730.8157 | 1794.6652 | 156.8971 |
| QD | 1729.6356 | 0.8243 | 15.9794 | 255.3408 | 1730.3328 | 1729.5616 | 1697.3776 | 1759.715 | 375.7582 |
| #Divergence_estimation_Clonal_Lineage_III |  |  |  |  |  |  |  |  |  |
|  | Mean | Stderr of mean | Stdev | Variance | Median | Geom. mean | Lower 95HPD | Upper 95HPD | ESS |
| Relaxed | 1756.2254 | 1.6702 | 17.9844 | 323.4379 | 1757.5543 | 1756.1331 | 1720.0277 | 1788.4566 | 115.9384 |
| Latorre | 1808.6322 | 1.0129 | 13.8883 | 192.8854 | 1809.8815 | 1808.5788 | 1781.2897 | 1834.8568 | 187.9903 |
| QD | 1795.4999 | 0.7965 | 12.3487 | 152.4911 | 1796.0612 | 1795.4574 | 1770.876 | 1819.0375 | 240.3418 |
| #Divergence_estimation_Clonal_Lineage_IV |  |  |  |  |  |  |  |  |  |
|  | Mean | Stderr of mean | Stdev | Variance | Median | Geom. mean | Lower 95HPD | Upper 95HPD | ESS |
| Relaxed | 1713.8149 | 1.5741 | 20.5187 | 421.0168 | 1715.334 | 1713.6916 | 1671.9086 | 1750.7509 | 169.8983 |
| Latorre | 1850.4954 | 0.7152 | 10.5547 | 111.4015 | 1851.3556 | 1850.4652 | 1828.617 | 1869.3638 | 217.7451 |
| QD | 1784.7554 | 0.6975 | 12.5339 | 157.0999 | 1785.4754 | 1784.7114 | 1759.7778 | 1808.4496 | 322.8878 |

**Table S5.** Comparison of Time of Most Recent Common Ancestor estimates with previous findings

| **Genetic Group** | **Estimate** | **Lower 95 HPD** | **Upper 95 HPD** |
| --- | --- | --- | --- |
| **Clonal Lineage II** | Original estimates* | 1744 | 1820 |
|  | This work (Latorre et al filtering) | 1731 | 1795 |
| **Clonal Lineage III** | Original estimates* | 1778 | 1847 |
|  | This work (Latorre et al filtering) | 1781 | 1835 |
| **Clonal Lineage IV** | Original estimates* | 1880 | 1916 |
|  | This work (Latorre et al filtering) | 1829 | 1869 |

\* Estimations from [8]

